# Neurofibromin is essential to maintain metabolic function and sustain life in the adult mouse

**DOI:** 10.1101/324061

**Authors:** Ashley N Turner, Maria S Johnson, Stephanie N Brosius, Brennan S. Yoder, Kevin Yang, Qinglin Yang, John F Moore, Qinglin Yang, John F Moore, J. Daniel Sharer, Daniel L Smith, Tim R Nagy, Trent R Schoeb, Bruce R Korf, Robert A Kesterson

**Author notes:** Corresponding Author: Robert A. Kesterson, PhD. Address: Department of Genetics, University of Alabama at Birmingham, Hugh Kaul Human Genetics Building, Rm 602, 720 20^th^ Street South, Birmingham, Alabama, USA 35294.

## Abstract

The consequences of pathogenic variants in the *NF1* gene can manifest in numerous tissues as a result of loss of neurofibromin protein function(s). A known function of *NF1* is negative regulation of p21ras signaling via a GTPase activating (Ras-GAP) domain. Besides modulation of Ras signaling as a tumor suppressor, other functions of this multi-domain protein are less clear. Biallelic inactivation of *NF1* leads to an embryonic lethal phenotype, while neurofibromin is expressed at varying levels in most tissues beyond developmental stages. Taking advantage of the mouse genetics toolkit, we established novel tamoxifen-inducible systemic knockout *Nf1* mouse models (C57BL/6) to gain a better understanding of the role of *Nf1* in the adult (3-4 months) mouse. Following inactivation of floxed *Nf1* alleles, adult *CAGGCre-ER*^*TM*^*;Nf1*^*4F/4F*^ mice lose function of *Nf1* systemically. Both male and female animals do not survive beyond 11 days post-tamoxifen induction and exhibit histological changes in multiple tissues. During this acute crisis, *CAGGCre-ER*^*TM*^*;Nf1*^*4F/4F*^ mice are not able to maintain body temperature or body mass, and expend all adipose tissue; however, they continue to consume food and absorb calories comparable to littermate-paired controls. Targeted metabolite analyses and indirect calorimetry studies revealed altered fat metabolism, amino acid metabolism and energy expenditure, with animals undergoing metabolic crisis and torpor-like states. Thermoneutral conditions accelerated the acute, lethal phenotype coincident with lower food intake. This study reveals that systemic loss of neurofibromin in the adult mouse induces metabolic dysfunction and lethality, thus highlighting potential functions of this multi-domain protein in addition to tumor suppression.

## Introduction

The *NF1* gene is considered a classical tumor suppressor and encodes neurofibromin, a multi-domain protein with the capacity to regulate several intracellular processes. The tumor suppressor function of neurofibromin is attributed to its negative regulation of p21ras signaling via a known GTPase-activating or GAP-related domain (GRD) (1–5). Mutations in *NF1* result in constitutive activation of Ras and downstream signaling cascades, including the Raf–MEK–ERK (MAPK) pathway and PI3K–Akt– mTOR pathway (6–8). Neurofibromin is also a positive regulator of adenylyl cyclase and involved in the cyclic AMP (cAMP) pathway (9, 10).

Much of our understanding of the mechanisms underlying the functional loss of *Nf1* come from mouse model studies. *NF1* deficiency in mice results in embryonic lethality due to heart abnormalities (11–13). Endeavors to create additional mouse models have largely focused around recapitulating manifestations seen in individuals affected with neurofibromatosis type 1 (NF1). NF1 is a common genetic disorder characterized by the occurrence of neurofibromas and café-au-lait macules, as well as many other features (14). NF1 is an autosomal dominant disorder with a near even divide between spontaneous and inherited mutations (14). Individuals with NF1 are at an increased risk for developing associated malignancies, and display an assortment of benign and malignant lesions (14).

Using conditional *Nf1* alleles in the mouse, numerous studies have explored the effects of *Nf1* loss in specific tissues. *Nf1* deletion in neurons results in abnormal development of the cerebral cortex and extensive reactive astrogliosis in the brain (15). Deletion of *Nf1* specifically in astrocytes in a *Nf1^+/−^* mouse leads to the development of optic nerve gliomas (16, 17). Additionally, *Nf1* loss in the Schwann cell lineage produces plexiform neurofibromas (13, 18, 19). Beyond tumorigenesis, *Nf1* loss in the early limb bud mesenchyme or muscle reveals abnormal skeletal muscle development, function, and metabolism in neonatal and adult mice (20–22).

To gain a better understanding of the role of *Nf1* in the adult mouse, *Nf1* was deleted in 3–4-month-old animals using the *Nf1*^*4Flox*^ conditional allele in combination with *CAGGCre-ER*^*TM*^, a tamoxifen-inducible ubiquitous *Cre* (13, 23). *Nf1* is lost only upon treatment of tamoxifen (TM). In adults, neurofibromin is expressed at varying levels in most tissues, suggesting an important role in multiple tissues. Our data show that induced loss of *Nf1* in adult animals is detrimental to metabolic function and viability.

## Materials and Methods

### Study approval

All experiments were conducted after approval of an animal protocol (APN130109835) by the Institutional Animal Care and Use Committee of the University of Alabama at Birmingham. All experiments were performed in accordance with the recommendations in the *Guide for the Care and Use of Laboratory Animals* published by the National Institutes of Health (NIH).

### Generation of mouse lines, breeding, and tamoxifen induction

Two *Nf1* alleles were utilized in this study as previously described (13). *Nf1*^*4Flox*^ (*Nf1*^*4F*^) is a conditional knockout allele of the *Nf1* gene with exon 4 flanked by loxP sites and *Nf1*^*Arg681\**^ is a nonsense mutation recapitulating human variant c.2041C>T;p.Arg681*.

Alleles were maintained on an inbred C57BL/6 background with animals group housed in a temperature- and humidity-controlled vivarium on a 12-hour dark-light cycle with *ad libitum* food (LabDiet NIH-31 Tac Auto/Irradiated; PMI Nutrition International, St. Louis, MO) and water access. To initiate a systemic knockout, a ubiquitously expressed tamoxifen inducible *CAGGCre-ER*^*TM*^ line was used to breed with *Nf1*^*4F/4F*^ and *Nf1*^*4F/Arg681\**^ mice (Figure S1A) (23). Male *CAGGCre-ER*^*TM*^;*Nf1*^*4F/4F*^ mice were crossed to *Nf1*^*4F/4F*^ or *Nf1*^*4F/Arg681\**^ females. Male and female animals were analyzed with Cre-negative *Nf1*^*4F/4F*^ or *Nf1*^*4F/Arg681\**^ littermates for controls. All animals had a background presence of Flp recombinase at the *Rosa26* locus.

Tamoxifen (TM, T5648; Sigma-Aldrich, St. Louis, MO) was dissolved in corn oil (C8267; Sigma-Aldrich, St. Louis, MO) at a concentration of 10 mg/mL. Three to four-month-old animals were injected intraperitoneally with vehicle or 6mg of TM/40g of body mass daily for 5 consecutive days (Fig.S1B). Initial induction of *Nf1*^*4F*^ allele loss was confirmed through genotyping followed by Western blotting. Animals were monitored for health and euthanatized if moribund per protocol (decrease greater than 20% of the initial body mass). Body mass was measured with an Ohaus Scout Pro SPE202 digital scale (Ohaus, Parsippany, NJ).

### Genotyping and Western blot analyses

Tail biopsies were obtained prior to and following TM induction to detect the presence of the converted null allele (*Nf1*^*Δ4*^). Genomic DNA was obtained by digesting tails in lysis buffer with proteinase K overnight, followed by phenol-chloroform extraction and ethanol precipitation. Polymerase chain reactions (PCRs) were set up with primer sets for detection of *Nf1*^*4F*^ allele, *Nf1*^*Δ4*^ allele, and *Cre* (Table S1). PCR products were analyzed on 1.5% agarose gels.

Tissue samples of adult brain were snap-frozen and homogenized by sonication in lysis buffer as previously described (13). Protein concentrations were determined using Pierce BCA Protein Assay Kit and 100μg total protein loaded per lane (Thermo Fisher Scientific, Waltham, MA). Western blot analysis was carried out as previously described (13). The membranes were immunoblotted with primary antibody at 4°C overnight, followed by incubation with secondary antibody (Table S2). Protein bands were visualized using Pierce ECL2 Western Blotting substrate (80196; Thermo Fisher Scientific, Waltham, MA) and the ChemiDoc XRS+ system (Bio-Rad, Hercules, CA). Equal loading was verified using α-tubulin.

### Histological and immunohistochemical analyses

Mice were intracardially perfused with 4% paraformaldehyde (PFA, w/v) in PBS. Tissues were dissected and post-fixed overnight, followed by paraffin embedding and sectioning. For histological analyses, tissues were cut into 5μm sections and stained with hematoxylin and eosin (H&E).

For immunohistochemical analysis, deparaffinized tissue underwent antigen retrieval and processing as previously described (24). Sections were incubated with antibody against Ki67 and then an HRP-conjugated goat anti-rabbit as previously described (Table S2) (24). Slides underwent Cy3 tyramide signal amplification and counterstaining with bisbenzimide as previously described (24). Images were acquired using a Nikon Eclipse TE-200 fluorescence microscope (Nikon, Minato, Tokyo, Japan) using Metamorph Imaging software (ver.6.1r6; Molecular Devices, San Jose, CA). Ten 40X fields were counted per animal, with a minimum of five animals counted per genotype.

### Temperature and body composition analyses

Rectal and surface temperature were measured using a MicroTherma 2T hand-held thermometer (TW2-107; ThermoWorks, American Fork, UT) with a rectal probe (RET 3; Physitemp Instruments, Clifton, New Jersey) and a FLIR T300 infrared (IR) imaging camera (FLIR Systems, Wilsonville, OR). IR images were analyzed using FLIR Tools software (ver.2.1; FLIR Systems, Wilsonville, OR).

Body composition was measured using the EchoMRI 3-in-1 composition analyzer Quantitative Magnetic Resonance (QMR) instrument (Echo Medical Systems, Houston, TX) as previously described (25).

### Hematological, clinical chemistry, and echocardiography analyses

Peripheral blood was collected in BD tubes (BD365963; Becton Dickinson and Company, San Jose, CA) by retro-orbital bleed under isoflurane micro-drop induction. The sample was kept on ice and submitted for complete blood count analysis to Charles River Research Animal Diagnostic Services (Wilmington, MA).

Peripheral blood was collected in BD tubes (BD365967; Becton Dickinson and Company, San Jose, CA) as described above. The clotted blood was centrifuged at 2,000 × g for 10 minutes at 4°C and the serum was transferred into a microcentrifuge tube, snap-frozen, and stored at −80°C until submitted for clinical chemistry analysis to Charles River Research Animal Diagnostic Services (Wilmington, MA).

Cardiac structure and function was assessed using the VEVO 770 echocardiograph system (FUJIFILM VisualSonics, Inc., Toronto, ON). Mice were anaesthetized with isoflurane inhalation with heart rate maintained at ~400 BPM and body temperature maintained at 37°C by placement on a heating pad. With a 35 MHz probe on long-axis and short axis M-mode images, wall thicknesses at diastole and systole were measured and analyzed using the manufacturer software package.

### Feeding and drinking analyses

Food/water intake was collected for individually housed mice that had undergone at least a one-week acclimation period from group to individual housing. These were measured daily as body mass described above.

### Bomb and indirect calorimetry analyses

Digestive efficiency was calculated by determining the energy content of food and feces by bomb calorimetry. Samples were weighed, dried, and energy content determined in triplicate with a Parr1261 calorimeter (Parr Instrument Company, Moline, IL). The average energy value was used for each sample and multiplied by the dry weight of sample to get energy in or out.

Indirect calorimetry measurements were conducted on individually housed mice that had undergone an acclimation period as noted above. Energy expenditure, respiratory exchange ratio (RER) and locomotor activity were acquired using a Labmaster indirect-calorimetry system (TSE Systems, Bad Homburg, Germany). Eight animals were individually housed in calorimetry chambers, fresh air was supplied at 0.42L/min, and O_2_ consumption and CO_2_ production were measured for 1 minute every 9 minutes for each. Total energy expenditure (TEE) was determined by calculating the average hourly energy expenditure over 22 hours multiplied by 24. Resting energy expenditure (REE) was determined as the average of the 3 lowest 18-minute periods. Non-resting energy expenditure (non-REE) was calculated as the difference between TEE and REE. Locomotor activity was determined with infrared beams for horizontal (x,y) activity. All animals were monitored daily for signs of distress. Animals were housed at room temperature (22°C) or thermoneutral conditions (30°C).

### Quantitative metabolite analyses

Peripheral blood was collected in BD tubes (BD365974; Becton Dickinson and Company, San Jose, CA) as described above. The sample was kept on ice and centrifuged at 2,000 × g for 15 minutes at 4°C. The clarified plasma was transferred to a microcentrifuge tube and stored at −80°C. Analysis of free amino acids in physiological samples was performed by high performance ion exchange liquid chromatography as described previously (26, 27). Plasma samples were first deproteinated with 10% 5-sulfosalycilic acid and then resolved via high performance ion exchange liquid chromatography using a discontinuous lithium chloride buffer gradient. Individual amino acids were detected by a post-column ninhydrin reaction and quantitated by comparing responses to a calibration curve.

Analysis of acylcarnitines in samples were performed by electrospray ionization-tandem mass spectrometry as described previously (28). Plasma samples were deproteinated by addition of methanol spiked with a mixture of eight internal standards. Acylcarnitines were converted to butyl esters and then subjected to electrospray ionization-tandem mass spectrometry; individual acylcarnitines were identified using a precursor ion scan and quantitated by comparison with the internal standards.

Organic acid analysis was performed on plasma and urine samples by gas chromatography-mass spectrometry as described (29). Urine was collected without intervention by allowing a mouse to urinate on wax paper outside of the cage. The voided urine was aspirated into a microcentrifuge tube and stored at −80°C. Plasma samples were initially deproteinated by adding an equal volume of 7% perchloric acid. After centrifugation, the protein-free supernatant was collected for analysis. Organic acids in the deproteinated plasma and urine samples were first oximated by incubation with hydroxylamine and then isolated by acidification and partitioning with 1:1 ethyl acetate/ethyl ether. The isolated organic acids were then derivatized with trimethylsilyltrifluoroacetamide/1% trimethylchlorosilane, and then subjected to gas chromatography-mass spectrometry. Organic acid-TMS derivatives were identified by comparing their characteristic fragmentation spectra to a mass spectra library and quantitated by comparing responses to a standard calibration curve.

### Statistical analysis

Statistical analyses were performed as indicated in the figure and table legends. Changes over time in continuous variables between groups including body mass, food intake, energy expenditure, and average respiratory exchange ratio were examined by repeated measures mixed models. A value of P<0.05 was considered statistically significant. Statistical analyses were performed using GraphPad Prism 7 software (GraphPad Software, Inc., La Jolla, CA) and SAS^®^ (ver.9.4.).

## Results

### Systemic induced Nf1 knockout leads to acute lethality

Following TM-induced inactivation of floxed *Nf1* alleles, *CAGGCre-ER*^*TM*^;*Nf1*^*4F/4F*^ adults lose function of *Nf1* systemically and are not able to survive beyond 11 days post-TM induction, showing a significant decrease in survival (Fig.1A). *CAGGCre-ER*^*TM*^;*Nf1*^*4F/4F*^ mice experience an acute phenotype and begin to show signs of hunched appearance and lordokyphosis prior to death. There was no observed phenotype or mortality with *Nf1*^*4F/4F*^ mice administered TM. Similarly, mice harboring the *Nf1*^*Arg681\**^ allele (nonsense mutation) with a single floxed *Nf1* allele (*CAGGCre-ER*^*TM*^;*Nf1*^*Arg681*/4F*^) fail to survive more than 11 days following inactivation of the floxed allele (Table S3).

**Figure 1.**
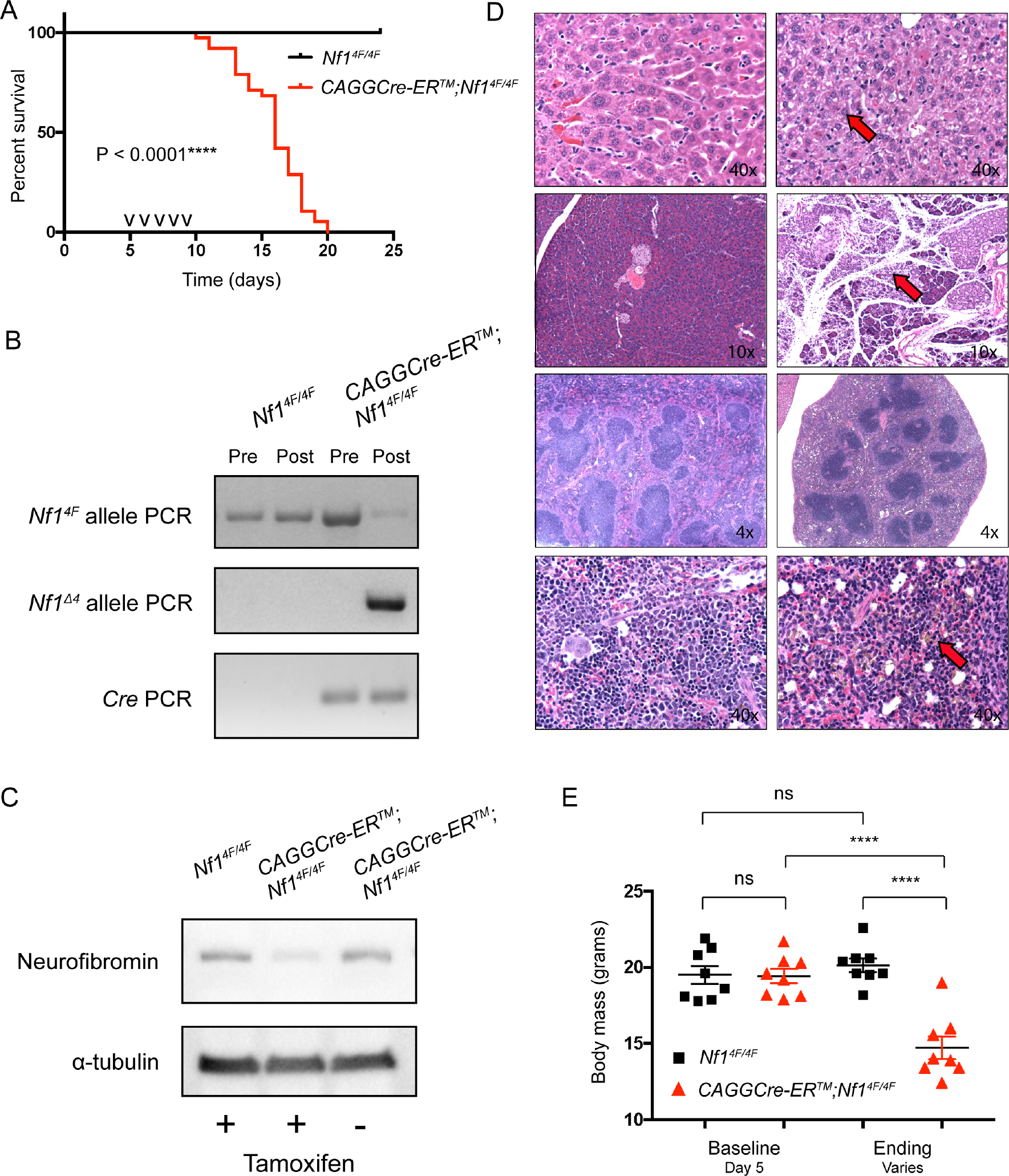
Tamoxifen-induced systemic knockout of *Nf1* in adult *CAGGCre-ER*^*TM*^*;Nf1*^*4F/4F*^ mice leads to acute, lethal phenotype with histological changes across multiple tissues and decreased body mass. (A) Kaplan-Meier curve showing no survival of *CAGGCre-ER*^*TM*^;*Nf1*^*4F/4F*^ mice (n=35, 13 male, 22 female) compared to *Nf1*^*4F/4F*^ mice (n=33, 14 male, 19 female) after TM induction by Log-rank Mantel-Cox test. Median survival 16 days, 6 days following TM induction. (B) Representative PCR analysis for a single animal’s tail DNA pre and post TM induction for *Nf1*^*4F/4F*^ and *CAGGCre-ER*^*TM*^;*Nf1*^*4F/4F*^ mice. Each PCR was set up with 50ng genomic DNA. The converted null *Nf1^Δ4^* allele is detected in *CAGGCre-ER*^*TM*^;*Nf1*^*4F/4F*^ animal following tamoxifen induction. (C) Representative Western blot of neurofibromin in whole brain lysate of indicated genotypes and TM treatment at day 12. α-tubulin is loading control.(D) Histological changes in *CAGGCre-ER*^*TM*^;*Nf1*^*4F/4F*^ mice compared to *Nf1*^*4F/4F*^ mice following TM induction at day 14. Representative H&E staining of liver (top panel), pancreas (second panel), and spleen (two bottom panels). *Nf1*^*4F/4F*^ is left side panels and *CAGGCre-ER*^*TM*^;*Nf1*^*4F/4F*^ is right side panels [n=4 per group (2 male, 2 female)]. Red arrowheads represent differences in liver (diffuse, mild individual cell hepatocellular degeneration and necrosis; hepatocytes are smaller); pancreas (patchy acinar atrophy); and spleen (smaller; severe loss lymphoid and hematopoietic elements; almost no erythropoiesis or myelopoiesis). (E) There is a significant difference in body mass between *Nf1*^*4F/4F*^ mice (n=8 female) and *CAGGCre-ER*^*TM*^;*Nf1*^*4F/4F*^ mice (n=8 female). ****P<0.0001, ns=not significant versus control by paired or unpaired t-test. Data are presented as mean±SEM. Tick mark indicates single TM injection. TM, tamoxifen.

Efficiency of the induced knockout of *Nf1* was assessed by PCR and Western blotting. Allele specific primer sets showed weak amplification of the *Nf1*^*4F*^ allele and strong amplification of the converted *Nf1*^*Δ4*^ null allele in *CAGGCre-ER*^*TM*^;*Nf1*^*4F/4F*^ mice after TM induction (Fig.1B). Since neurofibromin is highly expressed in nervous tissue, we used brain tissue as a proxy for TM efficiency. Western blotting for neurofibromin showed low levels in whole brain lysates of *CAGGCre-ER*^*TM*^;*Nf1*^*4F/4F*^ mice administered TM, indicating high efficiency of TM in the brain (Fig.1C).

While necropsies did not indicate a cause of death, histological analysis with H&E staining revealed histological changes in multiple tissues for *CAGGCre-ER*^*TM*^*;Nf1*^*4F/4F*^ mice compared to *Nf1*^*4F/4F*^ mice, including liver, pancreas, spleen, skin and bone marrow (Fig.1D; Fig.S2). No histological changes were noted in nervous tissues (Fig.S2). *CAGGCre-ER*^*TM*^;*Nf1*^*4F/4F*^ mice showed a significant decrease in body mass compared to their baseline body mass prior to TM and a significantly lower body mass compared to *Nf1*^*4F/4F*^ mice (Fig.1C). Body composition analysis revealed the difference of body mass to be attributed to a significantly lower amount of fat, lean and water mass for *CAGGCre-ER*^*TM*^;*Nf1*^*4F/4F*^ mice compared to *Nf1*^*4F/4F*^ mice (Fig.2). Gross dissections showed substantially lower fat mass in fat deposits across *CAGGCre-ER*^*TM*^;*Nf1*^*4F/4F*^ mice (Fig.2B). Body temperature analysis by rectal and IR thermometry revealed a decrease in the core body temperature of *CAGGCre-ER*^*TM*^;*Nf1*^*4F/4F*^ mice (Fig.S3).

**Figure 2.**
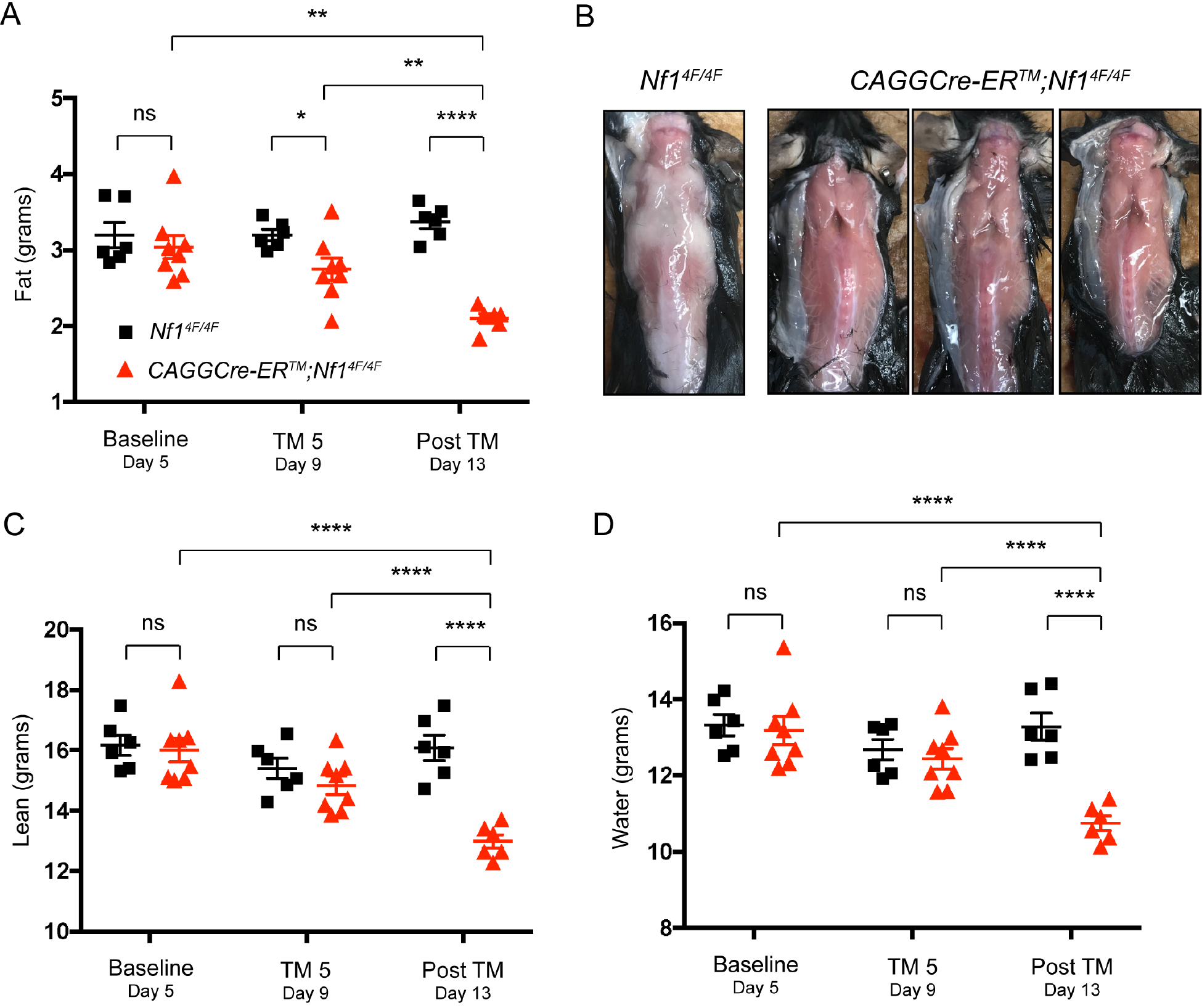
Body composition analysis with QMR reveals *CAGGCre-ER*^*TM*^;*Nf1*^*4F/4F*^ mice have lower mass for fat, lean, and water content following tamoxifen induction. (A) There is significantly lower body fat in *CAGGCre-ER*^*TM*^;*Nf1*^*4F/4F*^ (n=8 female) mice compared to *Nf1*^*4F/4F*^ mice (n=6 female) following TM induction. (B) Lower amount of fat visible in all fat deposits for *CAGGCre-ER*^*TM*^;*Nf1*^*4F/4F*^ mice compared to *Nf1*^*4F/4F*^ mice following TM induction at day 12, including scapular fat pads on dorsal side. (C-D) There is significantly lower lean (C) and water (D) in *CAGGCre-ER*^*TM*^;*Nf1*^*4F/4F*^ mice compared to *Nf1*^*4F/4F*^ mice following TM induction. *P<0.05, **P<0.01, ****P<0.0001, ns=not significant versus control by paired or unpaired t-test. Data are presented as mean±SEM. Tick mark indicates single TM injection. TM, tamoxifen.

To further examine the pathologies, complete blood count analysis, standard chemistry analysis, and echocardiography were performed. The red blood cell count was significantly higher in *CAGGCre-ER*^*TM*^;*Nf1*^*4F/4F*^ mice compared to *Nf1*^*4F/4F*^ mice following TM induction, whereas the white blood cell and lymphocyte counts were significantly lower (Fig.S4). Standard chemistry analysis revealed no significant differences across *CAGGCre-ER*^*TM*^;*Nf1*^*4F/4F*^ and *Nf1*^*4F/4F*^ mice with respect to triglycerides, free fatty acids, alanine aminotransferase, aspartate aminotransferase and glucose despite the observed changes in fat depots and histological differences in the liver and pancreas (Table 1). There were significant differences in circulating phosphorus and albumin levels (Table 1). Echocardiography revealed no significant difference in left ventricular anatomy or contractility. The systolic left ventricular internal diameter was significantly lower for *CAGGCre-ER*^*TM*^;*Nf1*^*4F/4F*^ compared to *Nf1*^*4F/4F*^ mice at day 12. Overall though, these results suggest that cardiac dysfunction was not the underlying cause of death in the *CAGGCre-ER*^*TM*^;*Nf1*^*4F/4F*^ mice (Table S4).

**Table 1.** Serum clinical chemistry levels in *CAGGCre-ER*^*TM*^;*Nf1*^*4F/4F*^ and *Nf1*^*4F/4F*^ mice following tamoxifen induction

|  |  | **Day 12** |  |  |
| --- | --- | --- | --- | --- |
|  |  | *Nf1<sup>4F/4F</sup>* (n=3) | *CAGGCre-ER<sup>TM</sup>;Nf1<sup>4F/4F</sup>* (n=3) | P-value |
|  |  | (Mean ± SEM) | (Mean ± SEM) |  |
| **Blood chemistry profiling** | **Units** |  |  |  |
| Cholesterol, total | mg/dL | 46.5 ± 3.86 | 48.33 ± 6.44 | 0.8056 |
| Triglycerides | mg/dL | 46.67 ± 5.78 | 38.67 ± 2.03 | 0.2618 |
| Free Fatty Acid (Non-esterified FA) | mmol/L | 0.30 ± 0.078 | 0.46 ± 0.06 | 0.1817 |
| Alanine Aminotransferase | U/L | 39.25 ± 9.47 | 49.67 ± 8.41 | 0.4669 |
| Aspartate Aminotransferase | U/L | 72.75 ± 14.15 | 294.3 ± 104 | 0.0544 |
| Alkaline Phosphatase | U/L | 85.5 ± 6.60 | 68 ± 12.58 | 0.2394 |
| Glucose | mg/dL | 164 ± 14.73 | 173 ± 55.75 | 0.8835 |
| **Phosphorus** | **mg/dL** | **8.33 ± 0.14** | **6.37 ± 0.27** | **0.0031\*\*** |
| Calcium, total | mg/dL | 8.57 ± 0.12 | 8.57 ± 0.03 | >0.9999 |
| Protein, total | g/dL | 4.33 ± 0.09 | 4.57 ± 0.09 | 0.1113 |
| **Albumin** | **g/dL** | **2.6 ± 0** | **2.83 ± 0.03** | **0.0004\*\*\*** |
| Urea Nitrogen | mg/dL | 12.75 ± 1.84 | 12.33 ± 0.33 | 0.8571 |
| Creatinine | mg/dL | 0.23 ± 0.03 | 0.23 ± 0.03 | 0.8457 |
| Bilirubin, total | mg/dL | 0.09 ± 0.02 | 0.09 ± 0.03 | 0.9808 |
| Sodium | mEq/L | 145.3 ± 2.40 | 141.7 ± 1.86 | 0.2938 |
| Potassium | mEq/L | 4.73 ± 0.17 | 4.6 ± 0.25 | 0.6815 |
| Chloride | mEq/L | 109.7 ± 2.19 | 106.3 ± 1.76 | 0.301 |

Data are presented as mean±SEM. Significant P-values are marked in bold with \*\*P<0.01 and \*\*\*P<0.001 versus control by unpaired t-test.

With observed histological changes for *CAGGCre-ER*^*TM*^;*Nf1*^*4F/4F*^ mice in metabolically active tissues, immunohistochemical analysis of Ki-67 staining was performed to assess proliferation. *CAGGCre-ER*^*TM*^;*Nf1*^*4F/4F*^ mice had a significantly lower proliferative index for intestine and skin compared to *Nf1*^*4F/4F*^ mice (Fig.S5A-F). No proliferative cells in the spleen of *CAGGCre-ER*^*TM*^;*Nf1*^*4F/4F*^ mice were found (Fig.S5G-I).

### Systemic induced Nf1 knockout has altered energy expenditure

With loss of Ki-67 staining in the intestine, this indicated a potential functional impairment that might be contributing to the phenotype in regards to food digestion, nutrient absorption, and metabolism. For metabolic analysis, mice were placed in an indirect-calorimetry system and monitored pre- and post-TM induction. *CAGGCre-ER*^*TM*^;*Nf1*^*4F/4F*^ mice showed significant decrease in body mass that was lower compared to *Nf1*^*4F/4F*^ mice following TM induction (Fig.3A). There were no significant differences in food consumption (Fig.3B), absorption of calories based on digestive efficiency, water consumption, activity, or energy balance (Fig.S6A-D). Both *CAGGCre-ER*^*TM*^;*Nf1*^*4F/4F*^ and *Nf1*^*4F/4F*^ mice showed a decrease in body mass and food intake after the first TM injection on day 6 (Fig.3A-B). This temporary energy imbalance cannot be fully attributed to the calories obtained from the corn oil vehicle, as this would only amount to 2.44–3.66 calories daily based on the range of injection volume administered.

**Figure 3.**
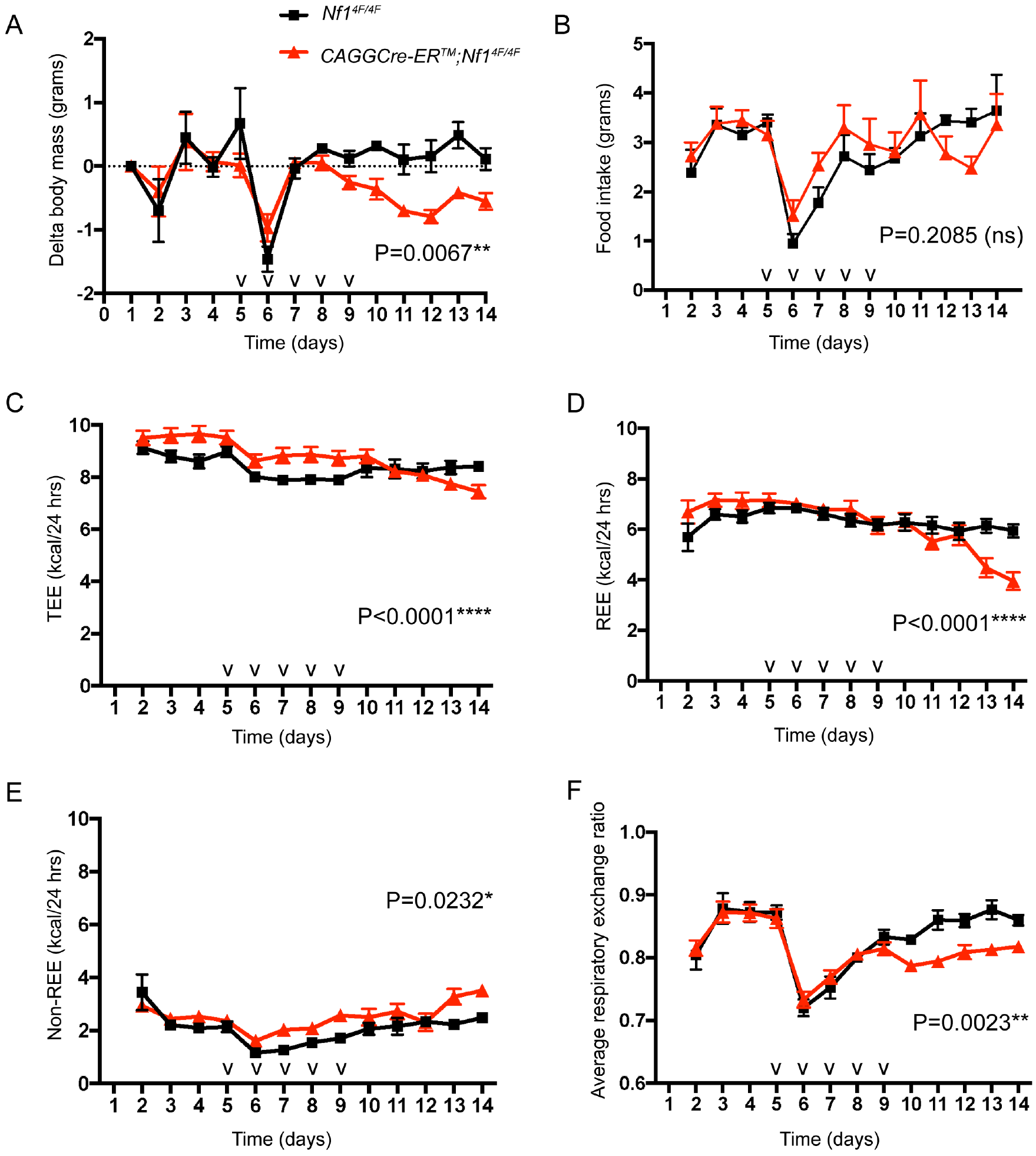
Metabolic characterization by indirect calorimetry reveals adult *CAGGCre-ER*^*TM*^*;Nf1*^*4F/4F*^ mice have altered total energy expenditure, resting energy expenditure, non-resting energy expenditure, and respiratory exchange ratio following tamoxifen induction. (A) Change in body mass is significantly lower for *CAGGCre-ER*^*TM*^;*Nf1*^*4F/4F*^ mice (n=8 female) compared to *Nf1*^*4F/4F*^ mice (n=6 female) following TM induction. (B) No significant difference detected with food intake. (C) TEE higher for *CAGGCre-ER*^*TM*^;*Nf1*^*4F/4F*^ mice during TM induction compared to *Nf1*^*4F/4F*^ mice. Following TM induction, TEE for *CAGGCre-ER*^*TM*^;*Nf1*^*4F/4F*^ mice decreases and is lower than *Nf1*^*4F/4F*^ mice. (D) Following TM induction, there is a decrease in REE for *CAGGCre-ER*^*TM*^;*Nf1*^*4F/4F*^ mice that is lower than *Nf1*^*4F/4F*^ mice. (E) After TM induction, non-REE increased in *CAGGCre-ER*^*TM*^;*Nf1*^*4F/4F*^ to be higher compared to *Nf1*^*4F/4F*^ mice. (F) Average RER decreased for *CAGGCre-ER*^*TM*^;*Nf1*^*4F/4F*^ mice after TM induction and is lower compared to *Nf1*^*4F/4F*^ mice. *P<0.05, **P<0.01, ****P<0.0001, ns=not significant versus control by repeated measures mixed models. Data are presented as mean±SEM. Tick mark indicates single TM injection. TM, tamoxifen.

There were significant differences between *CAGGCre-ER*^*TM*^;*Nf1*^*4F/4F*^ and *Nf1*^*4F/4F*^ mice for TEE, REE, non-REE, and average RER over time (Fig.3C-F). During TM induction, *CAGGCre-ER*^*TM*^;*Nf1*^*4F/4F*^ mice had a higher TEE and non-REE compared to *Nf1*^*4F/4F*^ mice (Fig.3C,E). Following TM induction, non-REE continued to be higher in *CAGGCre-ER*^*TM*^;*Nf1*^*4F/4F*^ mice compared to *Nf1*^*4F/4F*^ mice whereas TEE decreased in *CAGGCre-ER*^*TM*^;*Nf1*^*4F/4F*^ mice. Analysis of TEE difference across groups showed higher TEE during TM induction and lower TEE after TM for *CAGGCre-ER*^*TM*^;*Nf1*^*4F/4F*^ mice compared to *Nf1*^*4F/4F*^ mice (Fig.S6E). There was a significant decrease in REE and average RER after TM induction for *CAGGCre-ER*^*TM*^;*Nf1*^*4F/4F*^ mice that was lower than *Nf1*^*4F/4F*^ mice (Fig.3D,F). Examinations of individual animal’s energy expenditure and activity across the 24-hour monitoring period revealed the *CAGGCre-ER*^*TM*^;*Nf1*^*4F/4F*^ mice experience torpor-like states following TM induction (Fig.S7). Torpor has been traditionally defined as a temporary physiological state characterized by a controlled lowering of metabolic rate, body temperature, and physical activity, with a metabolic switch from consuming carbohydrates to consuming lipids usually during periods of low food availability (30, 31).

### Systemic induced Nf1 knockout has altered metabolism

Targeted metabolite analysis was performed to examine profiles for amino acids, lipids, and organic acids in the blood plasma and organic acids in the urine. Significant differences in the amino acid levels were observed in *CAGGCre-ER*^*TM*^;*Nf1*^*4F/4F*^ mice compared to *Nf1*^*4F/4F*^ mice on day 6 (following a single TM injection) and day 12 (three days post-TM induction). On day 6, *CAGGCre-ER*^*TM*^;*Nf1*^*4F/4F*^ mice have significantly lower levels of isoleucine and leucine and significantly higher level of ammonia (Table 2). By day 12, 31% (10/32) of amino acids are significantly lower in the *CAGGCre-ER*^*TM*^;*Nf1*^*4F/4F*^ mice compared to *Nf1*^*4F/4F*^ mice (Table 2). Taurine was the only amino acid that was significantly higher for *CAGGCre-ER*^*TM*^;*Nf1*^*4F/4F*^ mice. Only subtle differences in the plasma lipid levels were observed. On day 6, *CAGGCre-ER*^*TM*^;*Nf1*^*4F/4F*^ mice have significantly lower levels of C0 and C5, and significantly higher levels of C18-2 and C18-OH compared to *Nf1*^*4F/4F*^ mice (Table 3). The lipid profile changed slightly on day 12 with a significantly lower level of C0, and significantly higher levels of C5-1, C6, and C18-2 in *CAGGCre-ER*^*TM*^;*Nf1*^*4F/4F*^ mice compared to *Nf1*^*4F/4F*^ mice (Table 3). Few significant differences were noted in the organic acid levels in the urine, and no differences in the plasma on day 12. Glyceric and suberic acid, by-products of glycerol oxidation and fatty acid oxidation, were significantly higher in the urine of *CAGGCre-ER*^*TM*^;*Nf1*^*4F/4F*^ mice (Table 4). No differences were observed in ketone bodies with respect to 3-hydroxybutyric acid and total acetoacetic acid in urine or plasma.

**Table 2.** Plasma amino acid levels in *CAGGCre-ER*^*TM*^;*Nf1*^*4F/4F*^ and *Nf1*^*4F/4F*^ mice following tamoxifen induction

| (μmol/L) | Day 6 |  |  | Day 12 |  |  |
| --- | --- | --- | --- | --- | --- | --- |
|  | *Nf1<sup>4F/4F</sup>* (n=9)<br>(Mean ± SEM) | *CAGGCre-ER<sup>TM</sup>;Nf1<sup>4F/4F</sup>* (n=7)<br>(Mean ± SEM) | P-value | *Nf1<sup>4F/4F</sup>* (n=6)<br>(Mean ± SEM) | *CAGGCre-ER<sup>TM</sup>;Nf1<sup>4F/4F</sup>* (n=8)<br>(Mean ± SEM) | P-value |
| **Amino Acid Profiling** |  |  |  |  |  |  |
| Phosphoserine | 13.26 ± 2.28 | 12.03 ± 1.67 | 0.6868 | 16.63 ± 1.81 | 17.39 ± 1.47 | 0.7515 |
| **Taurine** | 685.4 ± 62.33 | 666.3 ± 65.13 | 0.8371 | **391.3 ± 24.37** | **527 ± 32.32** | **0.0126\*** |
| Phosphoethanolamine | 6.03 ± 1.44 | 5.86 ± 1.89 | 0.9441 | 9.75 ± 1.20 | 12.61 ± 1.39 | 0.1821 |
| Urea | 5541 ± 186.1 | 5064 ± 177 | 0.0911 | 5404 ± 391.8 | 5931 ± 599.9 | 0.5374 |
| **Aspartic acid** | 15.01 ± 0.45 | 15.9 ± 0.68 | 0.2758 | **20.42 ± 0.86** | **17.42 ± 0.79** | **0.0309\*** |
| Threonine | 138.6 ± 8.06 | 151.9 ± 14.27 | 0.4053 | 229.6 ± 27.12 | 188.3 ± 14.45 | 0.1677 |
| **Serine** | 117.9 ± 6.43 | 122 ± 11.21 | 0.7433 | **215.2 ± 20.71** | **128.8 ± 11.73** | **0.0023\*\*** |
| **Asparagine** | 56.37 ± 4.73 | 57.95 ± 4.34 | 0.8148 | **94.76 ± 10.74** | **57.8 ± 5.64** | **0.0063\*\*** |
| Glutamic acid | 21.74 ± 2.40 | 26.35 ± 3.41 | 0.2743 | 24.02 ± 5.29 | 31.14 ± 3.08 | 0.2352 |
| **Glutamine** | 792.7 ± 28.98 | 800.9 ± 32.09 | 0.8526 | **721 ± 52.36** | **517.6 ± 36.9** | **0.0083\*\*** |
| Alpha-aminoadipic acid | 5.79 ± 0.53 | 4.63 ± 1.25 | 0.3681 | 4.89 ± 0.56 | 10.81 ± 6.19 | 0.4731 |
| **Glycine** | 237.8 ± 16.2 | 249.5 ± 14.66 | 0.6111 | **364.5 ± 27.73** | **214.4 ± 20.81** | **0.0011\*\*** |
| **Alanine** | 451.9 ± 35.25 | 433.6 ± 34.93 | 0.7225 | **682.5 ± 69.41** | **427.6 ± 52.94** | **0.0133\*** |
| **Citrulline** | 51.88 ± 3.25 | 50.23 ± 4.6 | 0.7682 | **87.4 ± 9.35** | **53.88 ± 5.74** | **0.0077\*\*** |
| Alpha-aminobutyric acid | 11.62 ± 2.18 | 11.81 ± 1.52 | 0.9486 | 3.32 ± 0.49 | 14.32 ± 6.50 | 0.2149 |
| Valine | 150.2 ± 7.47 | 135.5 ± 10.7 | 0.2626 | 217.8 ± 22.32 | 190.8 ± 14.37 | 0.3074 |
| Cystine | 3.99 ± 1.26 | 3.15 ± 1.15 | 0.638 | 40.94 ± 13.45 | 15.4 ± 7.86 | 0.105 |
| **Methionine** | 56.01 ± 4.34 | 49.36 ± 4.33 | 0.304 | **116.4 ± 11.42** | **78.56 ± 9.13** | **0.0257\*** |
| **Isoleucine** | **73.22 ± 4.14** | **60.29 ± 3.85** | **0.0427\*** | 99.47 ± 9.52 | 82.66 ± 5.45 | 0.1253 |
| **Leucine** | **113.8 ± 6.22** | **86.14 ± 10.29** | **0.03\*** | 177.9 ± 19.14 | 145.2 ± 11.25 | 0.1421 |
| **Tyrosine** | 54.17 ± 3.21 | 52.93 ± 4.61 | 0.8232 | **114.5 ± 8.70** | **81.07 ± 7.38** | **0.0151\*** |
| Phenylalanine | 66.59 ± 2.30 | 61.57 ± 3.27 | 0.2167 | 91.91 ± 5.06 | 92.03 ± 5.40 | 0.9881 |
| Homocystine | 0.78 ± 0.39 | 0.99 ± 0.50 | 0.7463 | 3.76 ± 0.43 | 2.70 ± 0.48 | 0.1511 |
| **Ammonia** | **207.1 ± 14.96** | **274 ± 20.75** | **0.0178\*** | 243.2 ± 50.11 | 250.2 ± 29.11 | 0.8993 |
| Ornithine | 42.99 ± 2.43 | 49.85 ± 3.92 | 0.1418 | 86.89 ± 12.07 | 76.16 ± 14.57 | 0.6181 |
| Lysine | 267.3 ± 11.13 | 297.1 ± 19.99 | 0.1893 | 427.6 ± 32.38 | 395.7 ± 33.91 | 0.5381 |
| 1-methylhistidine | 6.19 ± 1.25 | 6.42 ± 1.37 | 0.9025 | 12.27 ± 0.86 | 10.78 ± 3.09 | 0.7181 |
| Histidine | 59.96 ± 1.84 | 54.43 ± 3.52 | 0.1593 | 77.82 ± 5.30 | 45.92 ± 12.64 | 0.071 |
| Tryptophan | 70.13 ± 6.48 | 63.38 ± 9.39 | 0.5507 | 74.51 ± 12.73 | 58.36 ± 16.25 | 0.4986 |
| **Arginine** | 86.25 ± 2.63 | 88.35 ± 5.14 | 0.703 | **127.8 ± 10.04** | **83.65 ± 13.04** | **0.0358\*** |
| Hydroxyproline | 21.77 ± 4.11 | 21.39 ± 7.73 | 0.9638 | 47.92 ± 8.42 | 35.69 ± 3.58 | 0.1517 |
| **Proline** | 71.94 ± 7.65 | 59.58 ± 7.19 | 0.2704 | **185.4 ± 28.3** | **103.1 ± 15.23** | **0.0169\*** |

Data are presented as mean±SEM. Significant P-values are marked in bold with \*P<0.05 and \*\*P<0.01 versus control by unpaired t-test.

**Table 3.** Plasma acylcarnitine levels in *CAGGCre-ER*^*TM*^;*Nf1*^*4F/4F*^ and *Nf1*^*4F/4F*^ mice following tamoxifen induction

| (μmol/L) | Day 6 |  |  | Day 12 |  |  |
| --- | --- | --- | --- | --- | --- | --- |
|  | *Nf1<sup>4F/4F</sup>* (n=9)<br>(Mean ± SEM) | *CAGGCre-ER<sup>TM</sup>;Nf1<sup>4F/4F</sup>* (n=7)<br>(Mean ± SEM) | P-value | *Nf1<sup>4F/4F</sup>* (n=6)<br>(Mean ± SEM) | *CAGGCre-ER<sup>TM</sup>;Nf1<sup>4F/4F</sup>* (n=8)<br>(Mean ± SEM) | P-value |
| **Lipid Profiling** |  |  |  |  |  |  |
| Acylcarnitines |  |  |  |  |  |  |
| **C0** | **53.66 ± 2.83** | **36.21 ± 3.77** | **0.0023\*\*** | **47.01 ± 4.80** | **34.79 ± 2.28** | **0.0344\*** |
| C2 | 40.75 ± 3.16 | 36.88 ± 2.60 | 0.3788 | 28.24 ± 6.72 | 23.27 ± 2.55 | 0.4783 |
| C3 | 1.62 ± 0.66 | 0.82 ± 0.13 | 0.3115 | 0.81 ± 0.08 | 0.70 ± 0.07 | 0.3535 |
| C4 | 0.60 ± 0.10 | 0.42 ± 0.06 | 0.216 | 0.51 ± 0.18 | 0.26 ± 0.03 | 0.1839 |
| **C5-1** | **0.08 ± 0.02** | **0.08 ± 0.03** | **0.9282** | **0.03 ± 0.002** | **0.07 ± 0.01** | **0.0137\*** |
| **C5** | **0.17 ± 0.02** | **0.11 ± 0.01** | **0.0464\*** | 0.10 ± 0.02 | 0.12 ± 0.01 | 0.391 |
| C4-OH | 0.50 ± 0.08 | 0.41 ± 0.08 | 0.4526 | 0.06 ± 0.01 | 0.12 ± 0.03 | 0.074 |
| **C6** | **0.15 ± 0.02** | **0.15 ± 0.04** | **0.8614** | **0.04 ± 0.01** | **0.10 ± 0.02** | **0.0201\*** |
| C6-OH | 0.06 ± 0.01 | 0.04 ± 0.004 | 0.1188 | 0.04 ± 0.01 | 0.04 ± 0.01 | 0.7565 |
| C8-1 | 0.15 ± 0.03 | 0.13 ± 0.04 | 0.6696 | 0.08 ± 0.02 | 0.08 ± 0.02 | 0.9364 |
| C8 | 0.10 ± 0.02 | 0.15 ± 0.03 | 0.1681 | 0.05 ± 0.01 | 0.09 ± 0.02 | 0.1332 |
| C3DC | 0.14 ± 0.02 | 0.16 ± 0.03 | 0.515 | 0.08 ± 0.01 | 0.11 ± 0.02 | 0.1565 |
| C10-1 | 0.11 ± 0.02 | 0.11 ± 0.04 | 0.9874 | 0.05 ± 0.02 | 0.10 ± 0.02 | 0.1564 |
| C10 | 0.10 ± 0.03 | 0.09 ± 0.02 | 0.6517 | 0.08 ± 0.01 | 0.08 ± 0.02 | 0.9167 |
| C4DC | 0.13 ± 0.02 | 0.14 ± 0.02 | 0.7314 | 0.06 ± 0.004 | 0.14 ± 0.05 | 0.1357 |
| C5DC | 0.20 ± 0.02 | 0.19 ± 0.02 | 0.7229 | 0.08 ± 0.03 | 0.06 ± 0.01 | 0.4284 |
| C12-1 | 0.10 ± 0.02 | 0.09 ± 0.02 | 0.8523 | 0.07 ± 0.04 | 0.09 ± 0.01 | 0.5562 |
| C12 | 0.18 ± 0.05 | 0.10 ± 0.02 | 0.1616 | 0.18 ± 0.10 | 0.10 ± 0.03 | 0.4405 |
| C6DC | 0.19 ± 0.06 | 0.29 ± 0.07 | 0.3125 | 0.13 ± 0.03 | 0.11 ± 0.01 | 0.5445 |
| C12-OH | 0.09 ± 0.01 | 0.1 ± 0.02 | 0.5893 | 0.11 ± 0.02 | 0.08 ± 0.01 | 0.2871 |
| C14-2 | 0.12 ± 0.02 | 0.12 ± 0.03 | 0.9921 | 0.10 ± 0.03 | 0.12 ± 0.04 | 0.7175 |
| C14-1 | 0.16 ± 0.05 | 0.16 ± 0.03 | 0.9427 | 0.1 ± 0.03 | 0.12 ± 0.02 | 0.5253 |
| C14 | 0.27 ± 0.06 | 0.31 ± 0.12 | 0.7529 | 0.10 ± 0.02 | 0.14 ± 0.05 | 0.4548 |
| C14-1-OH | 0.10 ± 0.02 | 0.09 ± 0.01 | 0.4873 | 0.08 ± 0.04 | 0.07 ± 0.02 | 0.8317 |
| C14-OH | 0.09 ± 0.02 | 0.12 ± 0.03 | 0.4061 | 0.10 ± 0.03 | 0.10 ± 0.02 | 0.8308 |
| C16-1 | 0.19 ± 0.05 | 0.16 ± 0.03 | 0.7277 | 0.07 ± 0.02 | 0.11 ± 0.03 | 0.2537 |
| C16 | 0.34 ± 0.04 | 0.34 ± 0.04 | 0.9559 | 0.19 ± 0.04 | 0.29 ± 0.05 | 0.1488 |
| C16-1-OH | 0.08 ± 0.01 | 0.07 ± 0.01 | 0.8635 | 0.08 ± 0.02 | 0.05 ± 0.01 | 0.1807 |
| C16-OH | 0.12 ± 0.02 | 0.08 ± 0.02 | 0.2871 | 0.07 ± 0.02 | 0.09 ± 0.02 | 0.5764 |
| **C18-2** | **0.17 ± 0.02** | **0.28 ± 0.04** | **0.0324\*** | **0.10 ± 0.02** | **0.20 ± 0.02** | **0.0012\*\*** |
| C18-1 | 0.30 ± 0.06 | 0.33 ± 0.06 | 0.7302 | 0.15 ± 0.04 | 0.25 ± 0.04 | 0.0946 |
| C18 | 0.24 ± 0.03 | 0.31 ± 0.10 | 0.4315 | 0.14 ± 0.03 | 0.18 ± 0.05 | 0.4697 |
| C18-2-OH | 0.06 ± 0.01 | 0.06 ± 0.02 | 0.9635 | 0.11 ± 0.04 | 0.11 ± 0.04 | 0.9632 |
| C18-1-OH | 0.08 ± 0.02 | 0.11 ± 0.02 | 0.4206 | 0.08 ± 0.02 | 0.07 ± 0.02 | 0.8175 |
| **C18-OH** | **0.06 ± 0.01** | **0.12 ± 0.02** | **0.0433\*** | 0.06 ± 0.01 | 0.09 ± 0.02 | 0.2219 |

Data are presented as mean±SEM. Significant P-values are marked in bold with \*P<0.05 and \*\*P<0.01 versus control by unpaired t-test.

**Table 4.** Urine and plasma organic acid levels in *CAGGCre-ER*^*TM*^;*Nf1*^*4F/4F*^ and *Nf1*^*4F/4F*^ mice following tamoxifen induction

| (μmol/L) | Day 12 |  |  |  |  |  |
| --- | --- | --- | --- | --- | --- | --- |
|  | Urine |  |  | Plasma |  |  |
|  | *Nf1<sup>4F/4F</sup>* (n=7)<br>(Mean ± SEM) | *CAGGCre-ER<sup>TM</sup>;Nf1<sup>4F/4F</sup>* (n=4)<br>(Mean ± SEM) | P-value | *Nf1<sup>4F/4F</sup>* (n=14)<br>(Mean ± SEM) | *CAGGCre-ER<sup>TM</sup>;Nf1<sup>4F/4F</sup>* (n=15)<br>(Mean ± SEM) | P-value |
| **Organic acid profiling** |  |  |  |  |  |  |
| Lactic acid | 171 ± 56.9 | 145.6 ± 65.12 | 0.7842 | 713.2 ± 99.53 | 780.9 ± 49.37 | 0.539 |
| Glycolic acid | 680.8 ± 177.2 | 625.5 ± 220.9 | 0.8522 |  |  |  |
| Glyoxylic acid | 208.4 ± 34.78 | 279.6 ± 82.84 | 0.3749 |  |  |  |
| Oxalic acid | 408.6 ± 67.17 | 462.7 ± 117.3 | 0.6735 |  |  |  |
| 3-hydroxypropionic acid | 26.98 ± 9.48 | 29.39 ± 14.06 | 0.8862 |  |  |  |
| Pyruvic acid | 38.26 ± 6.0 | 44.85 ± 8.96 | 0.5421 | 16.3 ± 2.27 | 17.48 ± 1.89 | 0.691 |
| 3-hydroxybutyric acid | 306.6 ± 177.4 | 64.55 ± 54.7 | 0.346 | 151.9 ± 49.84 | 216.5 ± 54.42 | 0.3913 |
| 2-hydroxyisovaleric acid | ND | ND | NA |  |  |  |
| 2-ketoisovaleric acid | 36.31 ± 12.16 | 52.76 ± 19.22 | 0.465 |  |  |  |
| Methylmalonic acid | ND | ND | NA |  |  |  |
| Total acetoacetic acid | 0.71 ± 0.71 | ND | 0.479 | ND | ND | NA |
| 2-keto-3-methylvaleric acid | 51.93 ± 12.46 | 79.99 ± 25.85 | 0.2942 |  |  |  |
| Ethylmalonic acid | 4.64 ± 0.81 | 5.75 ± 1.18 | 0.4434 |  |  |  |
| 2-ketoisocaproic acid | 76.36 ± 11.22 | 88.56 ± 17.81 | 0.5557 | 77.98 ± 8.48 | 91.23 ± 4.42 | 0.1692 |
| Succinic acid | 49.98 ± 14.59 | 46.73 ± 14.27 | 0.8875 | 4.40 ± 0.77 | 3.62 ± 0.50 | 0.3952 |
| **Glyceric acid** | **19.56 ± 7.18** | **89.7 ± 34.05** | **0.0266\*** |  |  |  |
| Fumaric acid | 5.92 ± 2.02 | 4.32 ± 1.92 | 0.6144 |  |  |  |
| Glutaric acid | 7 ± 2.45 | 6.06 ± 2.53 | 0.81 |  |  |  |
| 3-methylglutaric acid | 0.29 ± 0.21 | 0.64 ± 0.39 | 0.4073 |  |  |  |
| Succinylacetone | ND | ND | NA |  |  |  |
| Adipic acid | 3.94 ± 1.13 | 3.39 ± 1.41 | 0.7746 |  |  |  |
| 5-oxoproline | 196.6 ± 45.87 | 215.9 ± 57.22 | 0.8021 |  |  |  |
| 2-hydroxyglutaric acid | 68.45 ± 13.7 | 40.86 ± 7.97 | 0.1892 |  |  |  |
| 3-hydroxyglutaric acid | ND | ND | NA |  |  |  |
| 3-hydroxy-3-methylglutaric acid | ND | ND | NA |  |  |  |
| 2-ketoglutaric acid | 68 ± 13.54 | 60.04 ± 12.89 | 0.7078 |  |  |  |
| Para-hydroxy-phenylacetic acid | 37.59 ± 7.48 | 38.57 ± 9.84 | 0.9392 |  |  |  |
| **Suberic acid** | **0.47 ± 0.30** | **2.44 ± 0.10** | **0.0418\*** |  |  |  |
| Cis-Aconitic acid | 308.2 ± 59.71 | 158.2 ± 39.98 | 0.1146 |  |  |  |
| Orotic acid | ND | ND | NA |  |  |  |
| Citric acid | 432.5 ± 131.8 | 120.9 ± 28.9 | 0.1169 | ND | ND | NA |
| Methylcitric acid | ND | ND | NA |  |  |  |
| Sebacic acid | ND | ND | NA |  |  |  |
| Para-hydroxy-phenyllactic acid | 5.78 ± 1.55 | 6.7 ± 1.46 | 0.704 |  |  |  |
| Para-hydroxy-phenylpyruvic acid | 49.18 ± 9.38 | 64.82 ± 11.0 | 0.3239 |  |  |  |

Data are presented as mean±SEM. Significant P-values are marked in bold with \*P<0.05 versus control by unpaired t-test. ND, none detected. NA, not applicable.

### Impact of thermoneutral conditions on the acute, lethal phenotype

Indirect-calorimetry analysis was conducted in a second cohort at thermoneutrality (30°C). *CAGGCre-ER*^*TM*^;*Nf1*^*4F/4F*^ mice housed at 30°C have significantly lower survival compared to previous *CAGGCre-ER*^*TM*^;*Nf1*^*4F/4F*^ mice housed at room temperature (22°C) (Fig.4A). *CAGGCre-ER*^*TM*^;*Nf1*^*4F/4F*^ mice at 30°C are not able to survive beyond day 16 with a median survival of 13 days. There was no observable phenotype or mortality with *Nf1*^*4F/4F*^ mice administered TM at 30°C.

**Figure 4.**
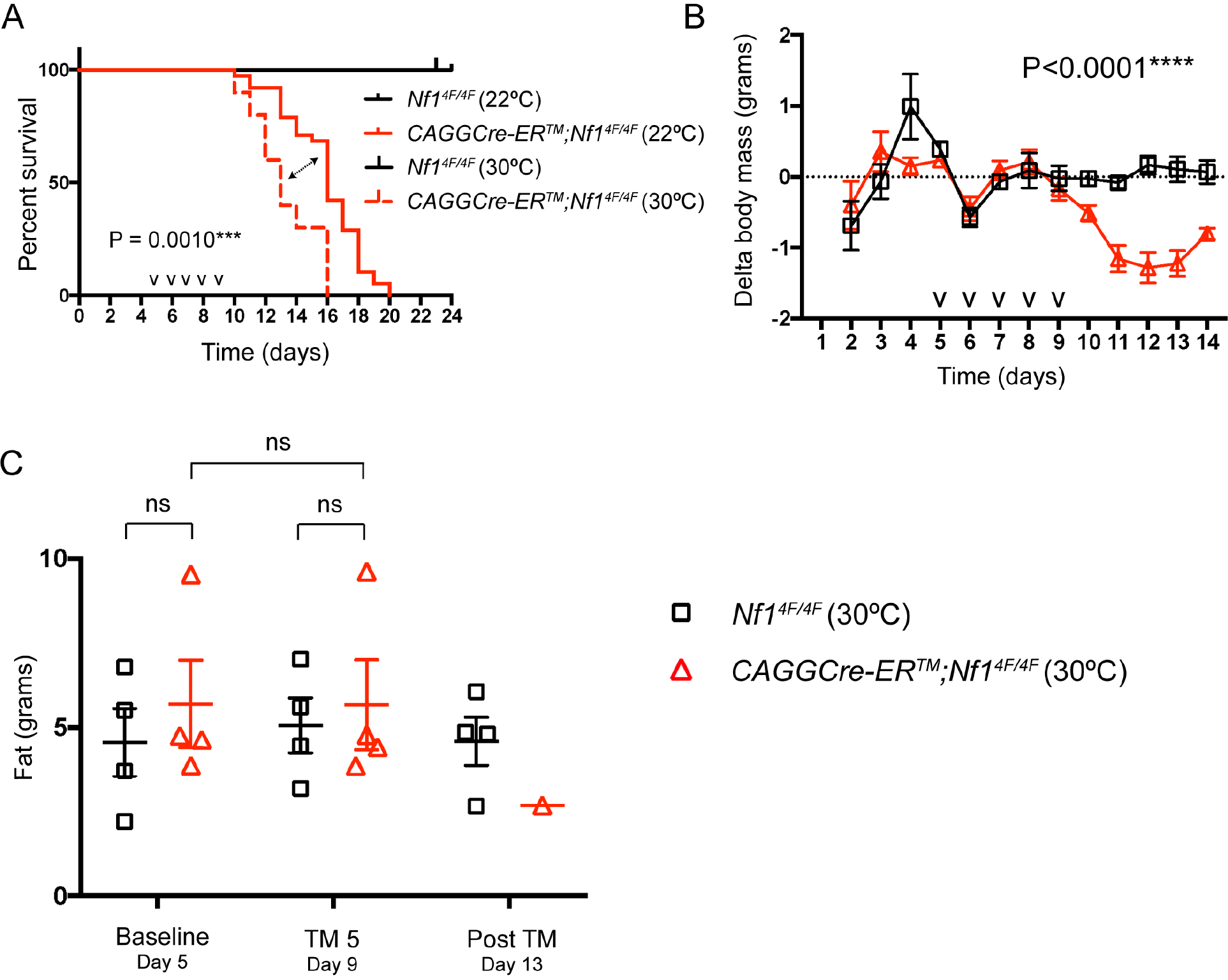
Thermoneutral conditions worsened the acute, lethal phenotype for adult *CAGGCre-ER*^*TM*^*;Nf1*^*4F/4F*^ mice. (A) Kaplan-Meier curve showing a significantly lower survival of *CAGGCre-ER*^*TM*^;*Nf1*^*4F/4F*^ mice at 30°C (n=10, 6 male, 4 female) compared to *CAGGCre-ER*^*TM*^;*Nf1*^*4F/4F*^ mice at 22°C (n=35, 13 male, 22 female) and *Nf1*^*4F/4F*^ at 30°C (n=10, 6 male, 4 female) and 22°C (n=33, 14 male, 19 female) after TM induction by Log-rank Mantel-Cox test. Median survival at thermoneutral conditions (30°C) is 13 days, 3 days following TM induction compared to 16 days, 6 days following TM induction at room temperature (22°C). A cross-sectional analysis was performed to include previous cohort at 22°C. (B) A lower body mass for *CAGGCre-ER*^*TM*^;*Nf1*^*4F/4F*^ mice compared to *Nf1*^*4F/4F*^ mice at 30°C. (C) There is a decrease in body fat in *CAGGCre-ER*^*TM*^;*Nf1*^*4F/4F*^ mice compared to *Nf1*^*4F/4F*^ at 30°C after TM induction. ***P<0.001, ****P<0.0001, ns=not significant versus control by paired or unpaired t-test or repeated measures mixed models. Data are presented as mean±SEM. Tick mark indicates single TM injection. TM, tamoxifen.

Changes in body mass and composition for *CAGGCre-ER*^*TM*^;*Nf1*^*4F/4F*^ mice were also observed at thermoneutrality. *CAGGCre-ER*^*TM*^;*Nf1*^*4F/4F*^ mice showed a significantly lower body mass compared to *Nf1*^*4F/4F*^ mice (Fig. 4B). Only a single *CAGGCre-ER*^*TM*^;*Nf1*^*4F/4F*^ mouse survived to day 13, and had lower body fat (Fig.4C). The body composition trends for fat, lean, and water mass were similar at both temperatures across time, being lower in *CAGGCre-ER*^*TM*^;*Nf1*^*4F/4F*^ mice compared to *Nf1*^*4F/4F*^ mice (Fig.S8A-C).

Body temperature analysis by rectal thermometry revealed a decrease in body temperature of *CAGGCre-ER*^*TM*^;*Nf1*^*4F/4F*^ mice that was lower than *Nf1*^*4F/4F*^ mice (Fig.5A). Thermoneutrality appears to have partially rescued the significantly lower rectal body temperature observed in *CAGGCre-ER*^*TM*^;*Nf1*^*4F/4F*^ mice at 22°C compared to 30°C (Fig.S9A). There were no torpor-like periods observed in *CAGGCre-ER*^*TM*^;*Nf1*^*4F/4F*^ mice at 30°C. There were no differences in activity between *CAGGCre-ER*^*TM*^;*Nf1*^*4F/4F*^ mice and *Nf1*^*4F/4F*^ mice at either 30°C or 22°C (Fig.S9B). *CAGGCre-ER*^*TM*^;*Nf1*^*4F/4F*^ and *Nf1*^*4F/4F*^ mice at 30°C were both in negative energy balance (Fig.S9C). *CAGGCre-ER*^*TM*^;*Nf1*^*4F/4F*^ mice had significantly lower food intake compared to *Nf1*^*4F/4F*^ mice at 30°C, in contrast to 22°C where no significant differences were observed (Fig.5B; Fig.S10B).

**Figure 5.**
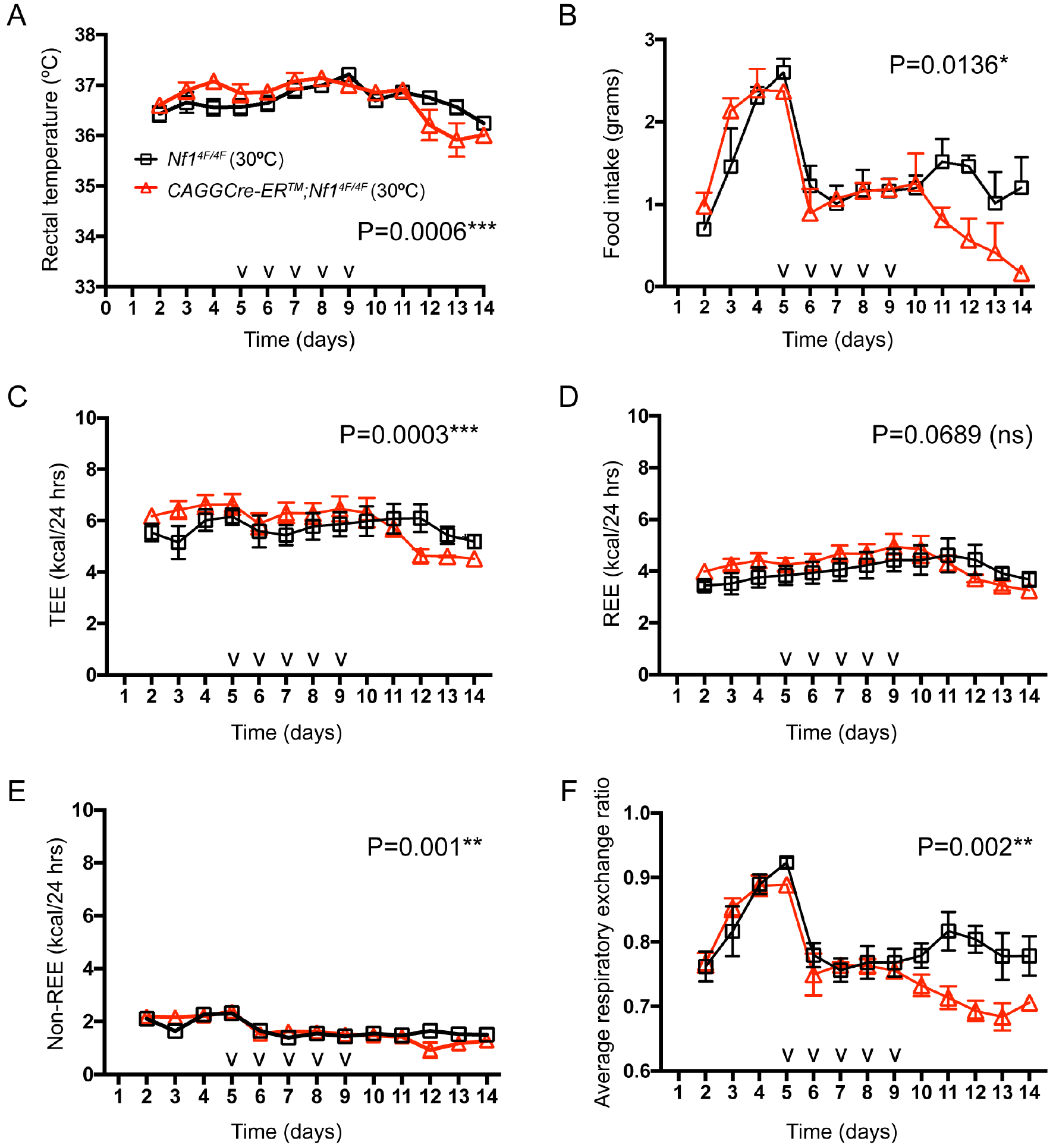
Significant differences observed with rectal temperature, food intake, total energy expenditure, non-resting energy expenditure, and average respiratory exchange ratio for adult *CAGGCre-ER*^*TM*^;*Nf1*^*4F/4F*^ mice following TM induction. (A) Significant difference in rectal body temperature between *CAGGCre-ER*^*TM*^;*Nf1*^*4F/4F*^ mice (n=10 with 6 male and 4 female) compared to *Nf1*^*4F/4F*^ mice at 30°C (n=10 with 6 male and 4 female). (B) There is significant difference of food intake between *CAGGCre-ER*^*TM*^;*Nf1*^*4F/4F*^ and *Nf1*^*4F/4F*^ (n=4 female per genotype). (C) TEE decreases for *CAGGCre-ER*^*TM*^;*Nf1*^*4F/4F*^ mice following TM induction and is lower than *Nf1*^*4F/4F*^ mice (n=4 female per genotype). (D) REE shows no significant difference at 30°C between *CAGGCre-ER*^*TM*^;*Nf1*^*4F/4F*^ and *Nf1*^*4F/4F*^ mice. (E-F) non-REE (E) and average RER (F) show significant differences at 30°C between *CAGGCre-ER*^*TM*^;*Nf1*^*4F/4F*^ and *Nf1*^*4F/4F*^ mice. *P<0.05, **P<0.01, ***P<0.001, ns=not significant versus control by repeated measures mixed models. Data are presented as mean±SEM. Tick mark indicates single TM injection. TM, tamoxifen.

At thermoneutrality, significant differences were observed between *CAGGCre-ER*^*TM*^;*Nf1*^*4F/4F*^ and *Nf1*^*4F/4F*^ mice for TEE, non-REE, and average RER (Fig.5C,E,F); however, there was no significant difference in REE (Fig.5D). These differences were similar to trends observed at 22°C, except for REE that decreased for *CAGGCre-ER*^*TM*^;*Nf1*^*4F/4F*^ mice at 22°C (Fig.S10C-F).

## Discussion

In this study, we employed a tamoxifen-inducible systemic knockout *Nf1* model to explore the role of *Nf1* in the adult mouse. First, we demonstrate that *Nf1* is essential for the viability of the adult mouse, and that neurofibromin is required throughout life.

Following systemic induced loss of *Nf1*, *CAGGCre-ER*^*TM*^;*Nf1*^*4F/4F*^ mice acutely lose body fat with no changes in food consumption or activity compared to *Nf1*^*4F/4F*^ mice. Necropsy revealed histological differences in numerous tissues of *CAGGCre-ER*^*TM*^;*Nf1*^*4F/4F*^ mice, but did not reveal the cause of death. These histologic differences indicate damage to metabolically active tissues. There was no evidence of infection or cardiovascular dysfunction that could explain the acute death. Recent studies show that NF1 patients have a high prevalence of being underweight with short stature accompanied by lower BMI (32–34). Further studies need to be conducted to assess body composition within Nf1 patients that could be underweight due to a decrease in body fat and a similar underlying metabolic condition as compared to this systemic knockout *Nf1* model.

Second, we discovered that the *CAGGCre-ER*^*TM*^;*Nf1*^*4F/4F*^ mice have altered energy expenditure and metabolism following systemic loss of *Nf1*. All evidence supports that the *CAGGCre-ER*^*TM*^;*Nf1*^*4F/4F*^ mice undergo an acute metabolic crisis that leads to torpor-like states. These torpor-like periods appear to be uncoupled from calorie restriction and reduced activity, suggesting this state is due to a metabolic response. *CAGGCre-ER*^*TM*^;*Nf1*^*4F/4F*^ mice have altered amino acid and fat metabolism. By-products from energy utilization cycles are not elevated, further suggest that they are also being utilized for energy in alternative pathways, such as ketone bodies. Recent studies with human cell lines and tissues have connected loss of neurofibromin with alterations in metabolism, specifically mitochondrial fatty acid metabolism and cellular respiration (20, 35). In addition, there appears to be an underlying metabolic phenotype in individuals with NF1. Adults with NF1 have a lower fasting blood glucose level compared with non-NF1 controls (36). Similarly, there are also repeated studies reporting a lower occurrence of diabetes mellitus in NF1 patients (37–39). A recent study found that NF1 patients have increased insulin sensitivity, lower levels of fasting blood glucose, and higher adiponectin levels (40). A study also observed excessive intake of saturated fatty acids and lipids in NF1 patients (41). The underlying reasons for each of these findings are still unclear, yet collectively suggest a role of metabolism and dietary intake affecting clinical manifestations of NF1. Our findings for the systemic induced loss of *Nf1* mouse model indicate fundamental changes in amino acid and fat metabolism that may explain these underlying NF1 clinical manifestations.

Additionally, the torpor-like states experienced by the *CAGGCre-ER*^*TM*^;*Nf1*^*4F/4F*^ mice do not appear to fit the traditional documented cases of torpor in mammals (30, 42). *CAGGCre-ER*^*TM*^;*Nf1*^*4F/4F*^ mice continue to consume food, absorb calories, and remain active comparable to *Nf1*^*4F/4F*^ mice. The findings suggest the acute, lethal phenotype is due to a hypermetabolic response. It is unclear why the *CAGGCre-ER*^*TM*^;*Nf1*^*4F/4F*^ mice do not attempt to compensate by increasing their dietary intake as they have *ad libitum* food access. This model could be used to further explore the disturbed signaling pathway(s) due to loss of *NF1* that provokes a metabolic-induced torpor-like state.

Third, thermoneutrality worsens the phenotype and lethality while partially rescuing the core body temperature and fully rescuing the hypometabolic, torpor-like states of *CAGGCre-ER*^*TM*^;*Nf1*^*4F/4F*^ mice. In our study, a higher dietary intake at room temperature appeared to be more protective than the decreased energy demand at thermoneutrality. This also suggests a sensitivity to thermal stress for the *CAGGCre-ER*^*TM*^;*Nf1*^*4F/4F*^ mice. Similar conclusions were reached investigating loss of *NF1* in *Drosophila melanogaster* with increased vulnerability to heat in *NF1* mutants (43).

This study illustrates the importance of considering a model’s physiological conditions. Most studies are conducted with cold-stressed mice due to the unique physiology of mice. This is an important variable of mouse physiology that should be considered in all studies trying to model human health and disease (44–46). The tamoxifen-inducible systemic knockout *Nf1* adult model is an example where room temperature versus thermoneutral conditions alters the phenotype in an unexpected way. Housing mice at thermoneutrality is a good practice when attempting to align mouse energy metabolism to human energy metabolism.

The tamoxifen-inducible knockout *Nf1* mouse model presented numerous challenges due to the nature of the systemic knockout phenotype. This has created the potential for confounding variables and insults leading to the phenotype and lethality. Additional studies need to assess the underlying molecular mechanisms of the phenotype. With the utilization of the *Nf1*^*4F*^ allele, the resulting *Nf1^Δ4^* allele is a very proximal null allele. Additional animal models with more distal and domain specific targeting could decouple and help elucidate neurofibromin function and the observed phenotype. Lastly, although Cre-negative control animals receiving TM did not show an observable phenotype, our studies do not rule out the possibility that loss of *Nf1* renders the mouse and tissues uniquely more sensitive to TM.

In summary, neurofibromin is essential in the adult mouse to maintain metabolic function, sustain life, and provides further insights into potential functions of this multi-domain protein. *CAGGCre-ER*^*TM*^;*Nf1*^*4F/4F*^ mice do not survive beyond 11 days post-tamoxifen induction and exhibit histological changes in multiple tissues. Targeted metabolite analysis and indirect calorimetry studies revealed altered fat and amino acid metabolism and energy expenditures, with animals undergoing metabolic crisis and torpor-like states. Thermoneutral housing suppressed the drop of body temperature and alleviated hypometabolic events, but worsened the acute, lethal phenotype. Further experiments will need to be undertaken to address mechanistic and translational questions still remaining. This study clearly shows that loss of *Nf1* is detrimental to metabolic function and viability in the adult mouse.

## Acknowledgments

The authors are grateful for the support of many organizations, programs, and sponsors to accomplish these studies. For assistance creating the mouse models, we would like to thank the UAB Transgenic & Genetically Engineered Models Systems (TGEMS) Core (supported by NIH awards P30CA13148, P30AR048311, P30DK074038, P30DK05336, and P60DK079626). We would like to acknowledge the UAB Animal Resources Program, UAB Comparative Pathology Laboratory, UAB Small Animal Phenotyping Core (supported by NIH awards P30DK056336, P30DK079626 and PAG050886A), and UAB Nutrition Obesity Research Center (supported by NIH award P30DK056336) for their technical assistance and support for these studies. This work was supported by the UAB Neurofibromatosis Program, through generous philanthropic gifts, and by the UAB Department of Genetics.

## Author Contributions

ANT was responsible for the experimental design, performing the experiments, data analysis and interpretation, and manuscript and figure preparation. MSJ carried out indirect calorimetry studies, designed the experiments, and analyzed the data. SNB assisted with immunohistochemistry staining. BSY assisted with feeding study. KY and QY carried out echocardiography study, designed the experiment, and analyzed the data. JFM and JDS carried out quantitative metabolite analyses. TRS performed pathological analysis of H&E stained tissue sections. QY, JDS, TRN, DLS, and BRK interpreted the data. RAK designed the experiments and interpreted the data. All authors discussed the results and implications and had final approval of the submitted and published versions.

## Supplementary Materials

Supplemental Tables (1-4); Supplemental Figures (1-10)

